# Gromov-Wasserstein optimal transport to align single-cell multi-omics data

**DOI:** 10.1101/2020.04.28.066787

**Authors:** Pinar Demetci, Rebecca Santorella, Björn Sandstede, William Stafford Noble, Ritambhara Singh

**Author notes:** Equal Contribution (Pinar Demetci, Rebecca Santorella).

## Abstract

Data integration of single-cell measurements is critical for understanding cell development and disease, but the lack of correspondence between different types of measurements makes such efforts challenging. Several unsupervised algorithms can align heterogeneous single-cell measurements in a shared space, enabling the creation of mappings between single cells in different data domains. However, these algorithms require hyperparameter tuning for high-quality alignments, which is difficult in an unsupervised setting without correspondence information for validation. We present Single-Cell alignment using Optimal Transport (SCOT), an unsupervised learning algorithm that uses Gromov Wasserstein-based optimal transport to align single-cell multi-omics datasets. We compare the alignment performance of SCOT with state-of-the-art algorithms on four simulated and two real-world datasets. SCOT performs on par with state-of-the-art methods but is faster and requires tuning fewer hyperparameters. Furthermore, we provide an algorithm for SCOT to use Gromov Wasserstein distance to guide the parameter selection. Thus, unlike previous methods, SCOT aligns well without using any orthogonal correspondence information to pick the hyperparameters. Our source code and scripts for replicating the results are available at https://github.com/rsinghlab/SCOT.

## 1 Introduction

Single-cell measurements provide a fine-grained view of the heterogeneous landscape of cells in a sample, revealing distinct subpopulations and their developmental and regulatory trajectories across time. The availability of single-cell measurements that capture various properties of the genome, such as gene expression, chromatin accessibility, DNA methylation, histone modifications, and chromatin 3D conformation, has increased the need for data integration methods capable of combining disparate data types.

Despite the importance of this task, the heterogeneity among single cells presents unique challenges. For example, due to technical limitations, it is hard to obtain multiple types of measurements from the same individual cell. Furthermore, when we measure different properties of a cell, we cannot a priori identify correspondences between features in the two domains. Accordingly, integrating two or more single-cell data modalities requires methods that do not rely on either common cells or features across the data types. This aspect prevents the application of some existing single-cell alignment methods to unsupervised settings because they require some correspondence information, either among the cells or the features [1–4]. For example, Seurat [4] requires correspondence information in the form of cells from similar biological state that are shared across the two datasets (known as *anchor points*). Cao *et al*. [5] have shown that such methods cannot perform good alignments under fully unsupervised settings.

Some approaches have tried to align datasets in an entirely unsupervised fashion. One of the earliest attempts, the joint Laplacian manifold alignment (JLMA) algorithm, constructs eigenvector projections based on local *k*-nearest neighbor graph Laplacians of the data [6]. The generalized unsupervised manifold alignment (GUMA) [7] algorithm seeks a 1–1 correspondence between two datasets based on a local geometry matching term. Liu *et al*. [8] showed that these methods do not perform well on the single-cell alignment task.

*Liu et al*. [8] proposed a manifold alignment algorithm based on the maximum mean discrepancy (MMD) measure, called MMD-MA, which can integrate different types of single-cell measurements. Another method, UnionCom [5], extends GUMA to perform unsupervised topological alignment. MMD-MA aims to match the global distributions of the datasets in a shared latent space, whereas UnionCom emphasizes learning both local and global alignments between the two distributions. Neither method requires any correspondence information either among samples or the features. The respective papers demonstrate state-of-the-art performance on simulated and real datasets. Although these results are encouraging, MMD-MA and UnionCom require that the user specify three and four hyperparameters, respectively. Selecting these hyperparameter values can be difficult and time-consuming in an unsupervised setting.

An emerging number of applications, including several in biology, are using optimal transport to learn a mapping between data distributions [9, 10]. Optimal transport finds the most cost-effective way to move data points from one domain to another. One way to think about it is as the problem of moving a pile of sand to fill in a hole through the least amount of work. Schiebinger *et al*. [11] use optimal transport to study how gene expression changes over time; they use regularized unbalanced optimal transport to compute differences in gene expression from one time point to the next. ImageAEOT [12] maps single-cell images to a common latent space through an autoencoder and then uses optimal transport to track cell trajectories. In related work, the same authors use autoencoders and optimal transport to learn transport maps among multiple domains [13]. However, the application of their method to single-cell datasets requires some form of supervision, like class labels, to preserve the underlying structure during transport.

The classic optimal transport problem requires datasets in the same metric space. Mémoli *et al*. [14] generalized optimal transport to the Gromov-Wasserstein distance, which compares metric spaces directly instead of comparing samples across spaces. In the natural language processing community, Alvarez *et al*. [10] used this approach to measure similarities between pairs of words across languages to compute the distances between languages. As far as we are aware, the only biological application of Gromov-Wasserstein optimal transport comes from [15], which uses it to reconstruct the spatial organization of cells from transcriptional profiles.

In this paper, we present Single-Cell alignment using Optimal Transport (SCOT), an unsupervised learning algorithm that uses Gromov-Wasserstein-based optimal transport to align single-cell multiomics datasets (presented schematically in Figure 1). Like UnionCom, SCOT aims to preserve local geometry when aligning single-cell data. SCOT achieves this by constructing a *k*-nearest neighbor (*k −*NN) graph for each dataset. SCOT then finds a probabilistic coupling between the samples of each dataset that minimizes the distance between the graph distance matrices produced by the *k*-NN graph. Finally, it uses the coupling matrix to project one single-cell dataset onto another through barycentric projection, thus aligning them. Unlike MMD-MA and UnionCom, SCOT requires tuning only two hyperparameters and is robust to the choice of one. We compare the alignment performance of SCOT with MMD-MA and UnionCom on four simulated and two real-world datasets. SCOT aligns all datasets as well as the state-of-the-art methods and scales well with increasing numbers of samples. Moreover, we demonstrate that the Gromov-Wasserstein distance can guide SCOT’s hyperparameter tuning in a fully unsupervised setting, when no orthogonal alignment information is available.

**Figure 1:**
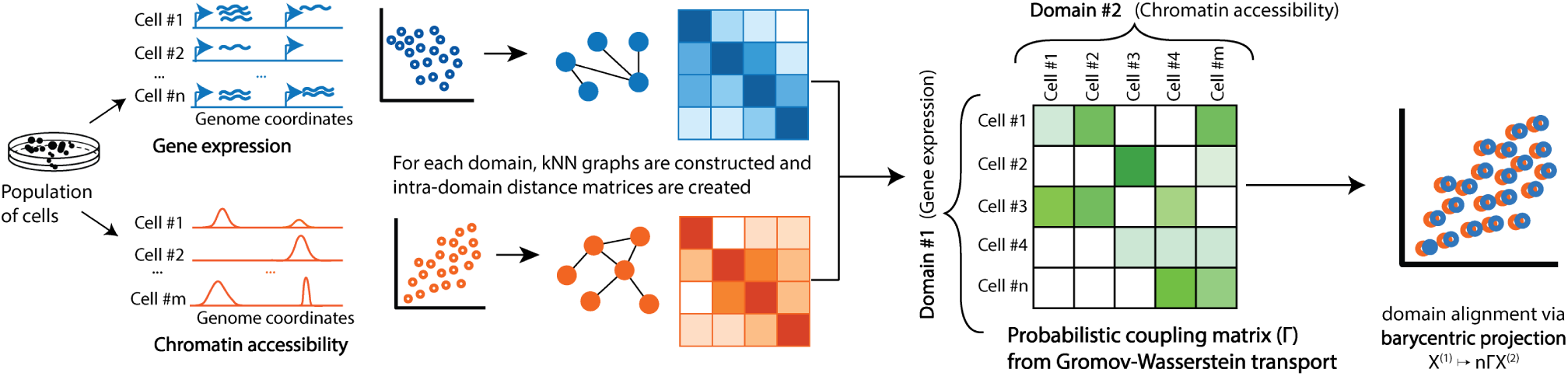
Schematic of SCOT alignment of single-cell multi-omics data. A population of cells is aliquoted for different single-cell sequencing assays to capture complementary aspects (e.g. gene expression and chromatin accessibility) of the molecular dynamics. SCOT constructs *k*-NN graphs based on sample-wise correlations, where vertices represent cells and finds a probabilistic coupling between the samples of each domain which minimizes the distance between the two intra-domain graph distance matrices. Barycentric projection uses this coupling matrix to project one domain onto another.

## 2 Methods

SCOT uses Gromov-Wasserstein optimal transport, which preserves local geometry when moving data points form one domain to another. The output of this transport problem is a matrix of probabilities that represent how likely it is that data points from one space correspond to data points in the other space. In this section, we introduce optimal transport followed by its extension to the Gromov-Wasserstein distance. Finally, we present the details of our SCOT algorithm.

We present the case for two datasets: 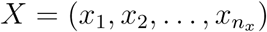 from *𝒳* and 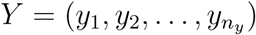 from 𝒴. The datasets have *n*_*x*_ and *n*_*y*_ points, respectively. We do not require any correspondence information but assume there is some underlying shared structure so that the datasets can be meaningfully aligned.

### Optimal transport

The Kantorovich optimal transport problem seeks to find a minimal cost mapping between two probability distributions [16]. Referring back to the problem of moving a sand pile to fill in a hole, Kantorovich optimal transport allows us to split the mass of a grain of sand instead of moving the whole grain; therefore, the mappings need not be 1—1. For probability measures *µ* and *ν* defined on *𝒳* and *𝒴*, respectively, this optimal transport problem finds a minimal coupling *π* that attains

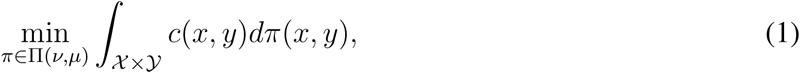

where *c*(*x, y*) is a cost function and П(*µ, ν*) is the set of couplings of *µ* and *ν* given by

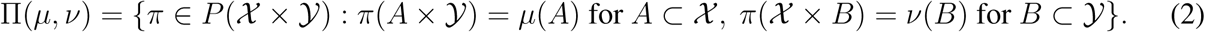

Intuitively, the cost function says how many resources it will take to move *x* to *y*, and the coupling *π* assigns a probability *π*(*x, y*) that *x* should be moved to *y*. When the spaces of interest are the same metric space with set *ℳ*, distance *d*, and cost function *c*(*x, y*) = *d*(*x, y*)^*p*^, the optimal transport distance (Equation 1) is equivalent to the *p−*th Wasserstein distance:

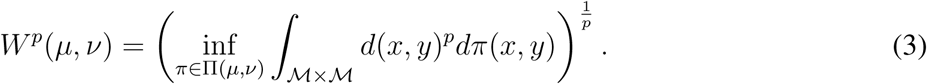

The Wasserstein distance measures the distances between probability distributions on a metric space and is commonly used in machine learning applications.

Since we align observed data points, we define the marginals as discrete empirical distributions:

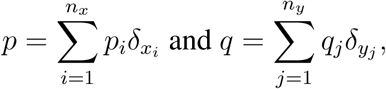

where 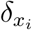 is the Dirac measure. Then, the cost function is given as a matrix 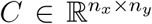, e.g. *C*_*ij*_ = ‖ *x*_*i*_ − *y*_*i*_ ‖, and the set of couplings are the matrices 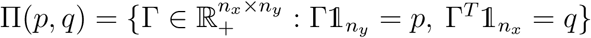. A discrete coupling Γ relates two measures *p* and *q*: each row Γ_*i*_ tells us how to split the mass of data point *x*_*i*_ onto the points *y*_*j*_ for *j* = 1, … *n*_*y*_, and the condition 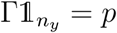 requires that the sum of each row Γ_*i*_ is equal to *p*_*i*_, the probability of sample *x*_*i*_. The discrete optimal transport problem finds a coupling that minimizes the cost of moving samples through the linear program:

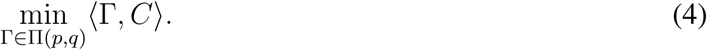

Although this problem can be solved with minimum cost flow solvers, it is usually regularized with entropy for more efficient optimization and empirically better results [17]. Entropy diffuses the optimal coupling, meaning that more masses will be split. Thus, the numerical optimal transport problem is

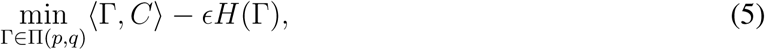

where *ϵ >* 0 and *H*(Γ) is the Shannon entropy defined as 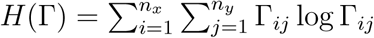.

Equation 5 is a strictly convex optimization problem, and for some unknownvectors 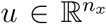 and 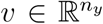 the solution has the form Γ^*∗*^ = diag(*u*)*K*diag(*v*), with *K* = exp, 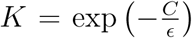 element-wise. This solution can be obtained efficiently via Sinkhorn’s algorithm, which iteratively computes

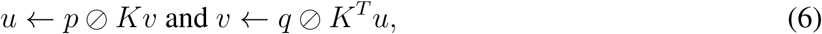

where ⊘ denotes element-wise division. This derivation immediately follows from solving the corresponding dual problem for Equation 5 [16].

### Gromov-Wasserstein distance

Classic optimal transport requires defining a cost function across domains, which can be difficult to implement when the domains are in different metric spaces. Gromov-Wasserstein distance extends optimal transport by comparing distances between samples rather than directly comparing the samples themselves [10]. We assume that we have metric measure spaces (𝒳, *d*_*x*_, *µ*) and (𝒴, *d*_*y*_, *ν*), where *d*_*x*_ and *d*_*y*_ are distances 𝒳 on and 𝒴, respectively [14]. Instead of defining a cost function between spaces, Gromov-Wasserstein uses the difference between pairwise distances. Given a cost function *L*: ℝ *×* ℝ *→* ℝ, the Gromov-Wasserstein distance between *µ* and *ν* is defined by

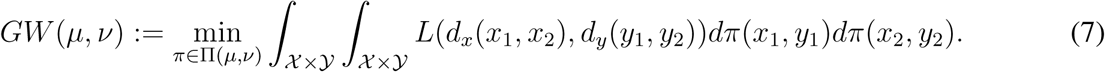

The main change from basic optimal transport (Equation 1) to Gromov-Wasserstein (Equation 7) is that we consider the effect of transporting pairs of points rather than single points. Intuitively, *L*(*d*_*x*_(*x*_1_, *x*_2_), *d*_*y*_(*y*_1_, *y*_2_)) captures how transporting *x*_1_ to *y*_1_ and *x*_2_ to *y*_2_ would distort the original distances between *x*_1_ and *x*_2_ and between *y*_1_ and *y*_2_. This change ensures that the optimal transport plan *π* will preserve some local geometry. In the case of 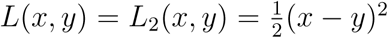, Gromov-Wasserstein is a distance on the space of metric measure spaces [14].

For the discrete case, we compute pairwise distance matrices *D*^*x*^ and *D*^*y*^ and the fourth order tensor 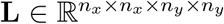, where 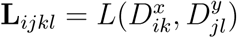. The discrete Gromov-Wasserstein problem is

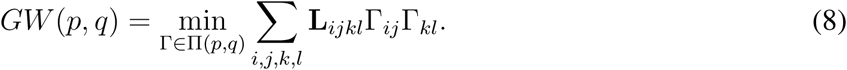

The summation can also be expressed as the inner product ⟨**L**(*D*^*x*^, *D*^*y*^) *⊗* Γ, Γ*⟩*. Equation 8 is now both non-linear and non-convex and involves operations on a fourth-order tensor, including the 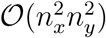 operation tensor product *L*(*D*^*x*^, *D*^*y*^) *⊗* Γ for a naive implementation. Peyre’ *et al*. show that for some choices of loss function this product can be computed in 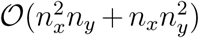 cost [18]. In particular, for the case *L* = *L*_2_, the inner product can be computed by

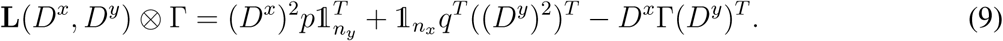

As in the classic optimal transport case, the coupling matrix can be efficiently computed for an entropically regularized optimization problem:

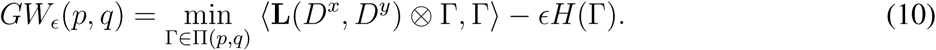

Larger values of *ϵ* lead to an easier optimization problem but also a denser coupling matrix, meaning that solutions will indicate significant correspondences between more data points. Smaller values of *ϵ* lead to sparser solutions, meaning that the coupling matrix is more likely to find the correct one-to-one correspondences for datasets where there are one-to-one correspondences. However, it also yields a harder (more non-convex) optimization problem [10].

Peyré *et al*. [18] propose using a projected gradient descent approach for optimization, where both the projection and the gradient are taken with respect to Kullback-Leibler divergence. These projections are computed via Sinkhorn iterations. Algorithm 1 in the supplement presents the algorithm for *L = L*_*2*_.

### Single-Cell alignment using Optimal Transport (SCOT)

Our method, SCOT, works as follows. First, we compute the pairwise distances on our data by using a geodesic distance as in [15]. To do this, we use the correlations between data points within each dataset to construct *k*-NN connectivity graphs. Then we compute the shortest path distance on the graph between each pair of nodes. We set the distance of any unconnected nodes to be the maximum (finite) distance in the graph and rescale the resulting distance matrix by dividing by the maximum distance. If *k* is the number of samples, then the *k*-NN graph is the complete graph, so the corresponding distance matrix is a matrix of all ones with zeros on the diagonal. In this case, the distance matrix does not provide information about the local geometry, so we recommend keeping *k* small relative to the number of samples to avoid this scenario. Our approach is robust to the choice of *k* (Supplementary Section 1.5)

Since we do not know the true distribution of the original datasets, we follow [10] and set *p* and *q* to be the uniform distributions on the data points. Then, we solve for the optimal coupling Γ which minimizes Equation 10. To implement this method, we use the Python Optimal Transport toolbox (https://pot.readthedocs.io/en/stable/) [19].

One of the advantages of using optimal transport is that we end up with a coupling matrix Γ with a probabilistic interpretation. The entries of the normalized row 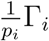 are the probabilitie-s that the fixed data point *x*_*i*_ corresponds to each *y*_*j*_. However, to use the correspondence metrics previously used in the field to evaluate the alignment, we need to project the two datasets into the same space. The Procrustes approach proposed in [10] does not generalize to datasets with different feature and sample dimensions, so we use a barycentric projection:

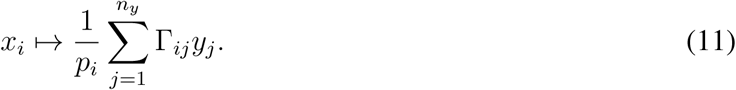

### Alternative Unsupervised Alignment Procedure

In the description of SCOT, the number *k* for nearest neighbors and the entropy weight *E* are hyperparameters. One way to set these hyperparameters for optimal alignment is to use some orthogonal correspondence information to select the best alignment either directly [5, 8] or by performing cross-validation [20]. This selection strategy is problematic for truly unsupervised setting, where no correspondence information is available a priori. As a solution, we provide an alternative procedure to learn reasonable alignments based on tracking the Gromov-Wasserstein distance (Equation 8). This procedure is based on our observation that the Gromov-Wasserstein distance serves as a proxy for measuring alignment quality (see Supplementary Figure S5). In this procedure, we alternate between optimizing *ϵ* and *k* to minimize the Gromov-Wasserstein distance between the domains (detailed in Algorithm 2 in Supplementary Materials). Although the lowest Gromov-Wasserstein distance is not always the best alignment, it consistently appears to be one of the better alignments.

## 3 Experimental Setup

### Simulated datasets

We follow Liu *et al*. [8] and benchmark SCOT on three different simulations^1^. All three simulations contain two domains with 300 samples that have been non-linearly projected to 1000- and 2000-dimensional feature spaces, respectively. The three simulations are a bifurcation, a Swiss roll, and a circular frustum (Supplementary Figure S1) with points belonging to three different groups. In addition to these three previously existing simulations, we use Splatter [21] to create simulated single-cell RNA sequencing count data, which we call synthetic RNA-seq. We generate 5000 cells with 1000 genes from three cell groups and reduce the count matrix to the five genes with the highest variances. This count matrix is randomly mapped into two new domains with dimensions *p*_*1*_ = 50 and *p*_*2*_ = 500 by multiplying it with two randomly generated matrices, resulting in data with dimensions 5000*×*50 and 5000×500.

All four datasets were simulated with 1—1 sample-wise correspondences, which are solely used for evaluating model performance. Each domain is projected to a different dimension, so there is no featurewise correspondence either. In all simulations, we Z-score normalize the features before running the alignment algorithms as in [8].

### Single-cell multi-omics datasets

We use two sets of single-cell multi-omics data to demonstrate the applicability of our model to real datasets. Both datasets are generated by co-assays; thus, we have known cell-level correspondence information for benchmarking. The first dataset is generated using the scGEM assay [22], which simultaneously profiles gene expression and DNA methylation. The dataset (Sequence Read Archive accession SRP077853) is derived from human somatic cell samples undergoing conversion to induced pluripotent stem cells (iPSCs). This dataset was also used by Cao *et al*. [5] to demonstrate the performance of their UnionCom algorithm. The data dimensions are 177*×*34 for the gene expression data and 177*×*27 for the chromatin accessibility data.

The second dataset is generated by the SNAREseq assay [23], which links chromatin accessibility with gene expression. The data (Gene Expression Omnibus accession GSE126074) is derived from a mixture of human cell lines: BJ, H1, K562, and GM12878. We pre-process the datasets following Chen *et al*. [23], as follows. We reduce data sparsity and noise in the ATAC-seq data by performing dimensionality reduction using the topic modeling framework cisTopic [24]. The dimensions of the RNA-seq data were reduced using PCA. The resulting input matrices for the SNARE-seq data were of size 1047×19 and 1047×10 for ATAC-seq and RNA-seq, respectively. We unit normalize all real datasets as done in [20].

### Evaluation metrics

We compare SCOT with the two state-of-the-art unsupervised single-cell alignment methods MMD-MA [8] and UnionCom [5]. None of these methods use any correspondence information for aligning the datasets. However, all datasets have 1–1 sample-level correspondence information, which we use to quantify the alignment performance through the “fraction of samples closer than the true match” (FOSCTTM) metric introduced by Liu *et al*. [8]. For each domain, we compute the Euclidean distances between a fixed sample point and all the data points in the other domain. Next, we use these distances to compute the fraction of samples that are closer to the fixed sample than its true match. Finally, we average these values for all the samples in both domains. For perfect alignment, all samples would be closest to their true match, yielding an average FOSCTTM of zero. Therefore, a lower average FOSCTTM corresponds to better alignment performance.

Since all the datasets have group-specific (simulations) or cell-type-specific (real experiments) labels, we also adopt the metric used by *Cao et al*. [5] called “label transfer accuracy” to assess the quality of the cell label assignment. It measures the ability to correctly transfer sample labels from one domain to another based on their neighborhood in the aligned domain. As described in [5], we train *a k*-nearest neighbor classifier on one of the domains and predict the sample labels in the other domain. The label transfer accuracy is the proportion of correctly predicted labels, so it ranges from 0 to 1, and higher values indicate good performance. We apply this metric to alignments selected by the FOSCTTM measure.

### Hyperparameter tuning

We run each method over a grid of hyperparameters and select the setting that yields the lowest average FOSCTTM. For SCOT, the grid covers the regularization weight ϵ ∈ {0.0001, 0.0005, 0.001, 0.005, …, 0.1} and number of neighbors 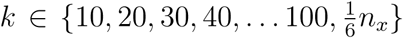. We observe empirically that going above 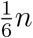 for *k* does not yield any improvement in alignment.

We pick the hyperparameters for MMD-MA and UnionCom based on the default values and recommended ranges. MMD-MA has three hyperparameters: weights *λ*_1_, *λ*_2_ ∈ {10^*−*3^, 10^*−*4^, 10^*−*5^, 10^*−*6^, 10^*−*7^} for the terms in the optimization problem and the dimensionality *p ∈* {4, 5, 6, 16, 32, 64} of the embed-ding space. UnionCom requires the user to specify four hyperparameters: the number *kmax ∈* {40, 100}of maximum number of neighbors in the graph,the dimensionality *p ∈* {4, 5, 6, 16, 32, 64} of the embedding space, the trade-off parameter *β ∈* {0.1, 1, 10, 15, 20} for the embedding, and a regularization coefficient *ρ ∈* {0, 5, 10, 15, 20}. We select the embedding dimension *p ∈* {16, 32, 64} around the default value of 32 set by UnionCom but also add *p ∈* {4, 5, 6} to match the recommended values for MMD-MA. We keep the hyperparameter search space size approximately consistent across the three methods. For each dataset, we present alignment and runtime results for the best performing hyperparameters.

Furthermore, we consider the scenario where correspondence information is unavailable to pick the optimal hyperparameters. For SCOT, we apply the alternative unsupervised alignment algorithm (Algorithm 2 in Supplementary Materials) to align all the datasets. Since MMD-MA and UnionCom do not provide a hyperparameter selection strategy, we rely on the default hyperparameters; we use Union-Com’s provided default parameters of *kmax* = 40, *p* = 32, *ρ* = 10, and *β* = 1, and the center values of MMD-MA’s recommended range: *p* = 5, *λ*_1_ = 10^*−*5^, and *λ*_2_ = 10^*−*5^. We also present the alignment results for all three methods in this fully unsupervised setting.

## 4 Results

### SCOT successfully aligns the simulated datasets

We first compare SCOT’s performance with MMD-MA and UnionCom for the four simulation datasets. In this experiment, we select the best performing hyperparameters for each method using the tuning process described in the previous section. In Figure 2, we sort and plot the FOSCTTM score for each sample for the simulations from [8], as well as the synthetic RNA-seq count data from Splatter [21]. Overall, we observe that SCOT consistently achieves one of the lowest average FOSCTTM scores, thereby demonstrating its ability to recover the correct correspondences. We also report the label transfer accuracy results (Table 4) when the first domain is used to train a classifier to predict the labels in the second domain. We observe that SCOT consistently yields high label transfer accuracy scores, indicating that samples are correctly mapped to their assigned groups.

**Figure 2:**
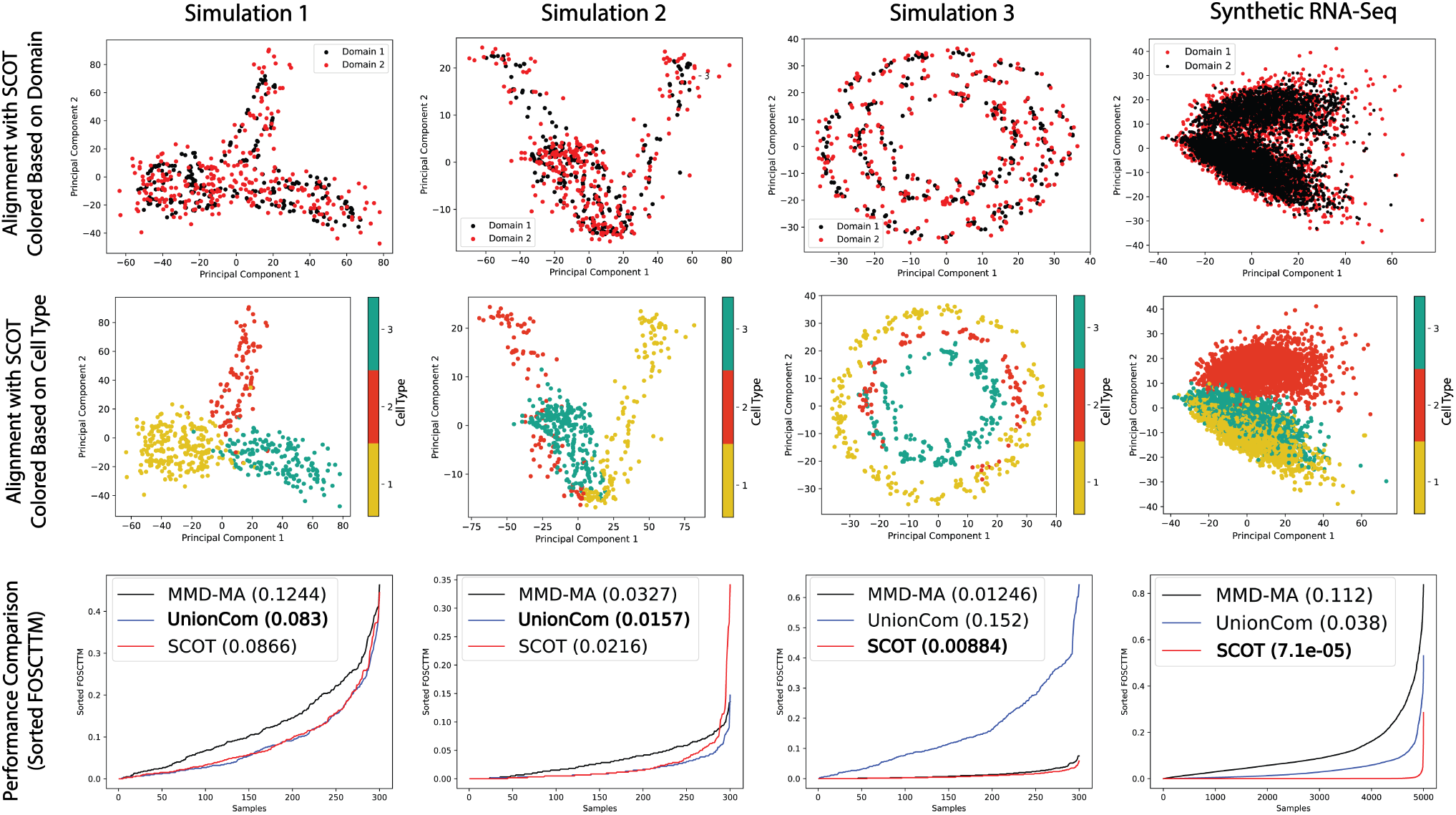
Aligning simulated datasets. Each column presents a different simulation. **Top:** our alignment colored by domain (plotted in 2D using PCA). **Middle:** our alignment colored by group. **Bottom:** sorted “fraction of samples closer than the true match” (FOSCTTM) for MMD-MA, UnionCom, and SCOT to visualize the distribution across samples with the average FOSCTTM values in the legend.

### SCOT gives state-of-the-art performance for single-cell multi-omics alignment

Next, we apply our method to real single-cell sequencing data. Overall, SCOT gives the lowest average FOSCTTM measure in comparison to MMD-MA and UnionCom (Figure 3, last column) and recovers accurate 1–1 corre-spondences in single-cell datasets. For the scGEM data, we report label transfer accuracy using the DNA methylation domain for predicting the cell-type labels in the gene expression domain. For the SNARE-seq dataset, we use the gene expression domain for predicting cell labels in the chromatin accessibility domain. SCOT yields the best label transfer accuracy result on SNAREseq dataset and performs comparably to the other methods for scGEM (Table 4.)

**Figure 3:**
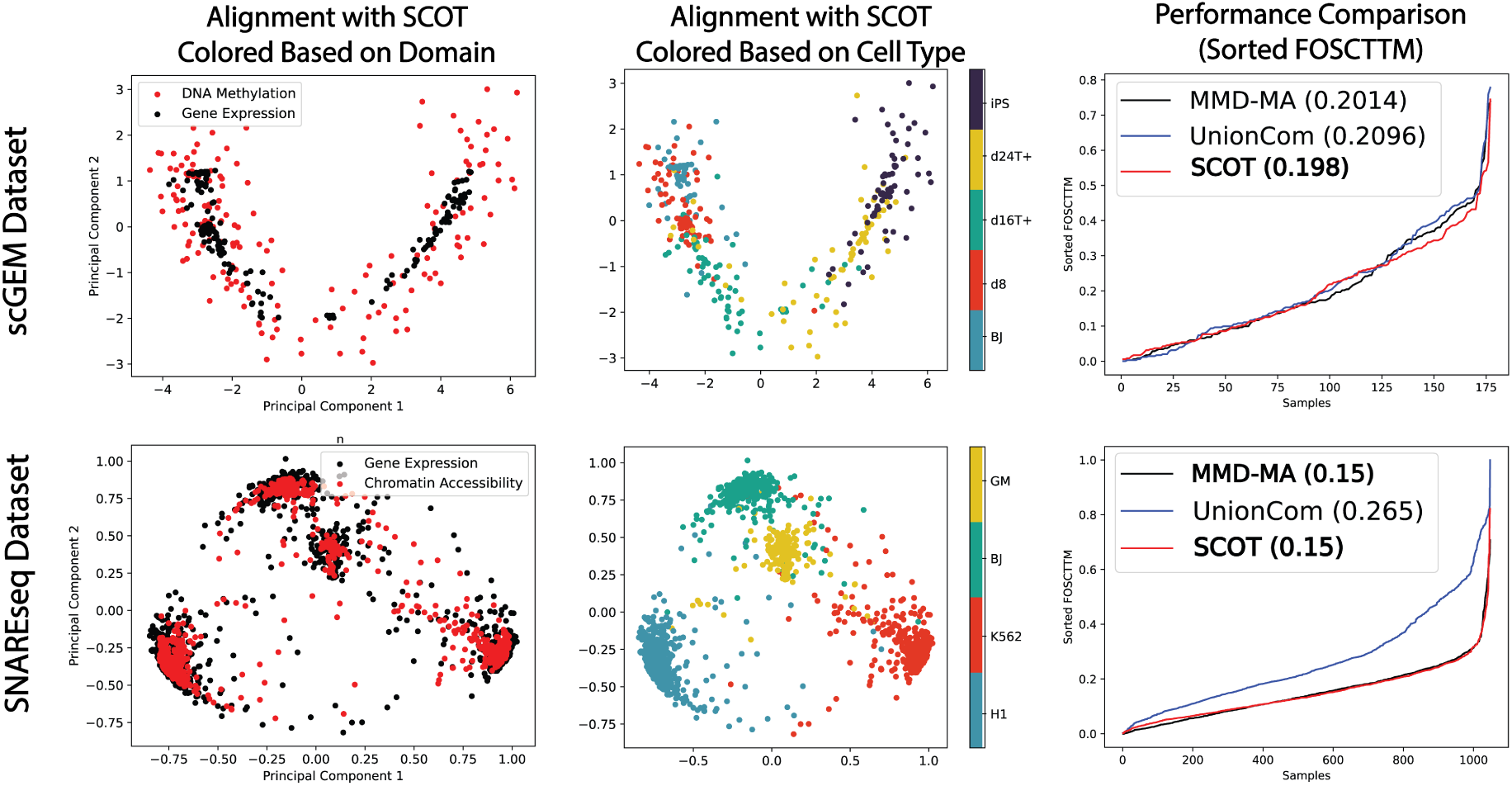
Aligning real world single-cell sequencing dataset. Each row presents a different real-word single-cell sequencing dataset. **Left:** our alignment colored based by domain (plotted in 2D using PCA). **Middle:** our alignment colored by cell-type. **Right:** sorted “fraction of samples closer than the true match” (FOSCTTM) for MMD-MA, UnionCom, and SCOT order to visualize the distribution across samples with the average FOSCTTM values in the legend.

While MMD-MA and UnionCom project both datasets to a shared low-dimensional space, SCOT projects one dataset onto the other. We project SCOT in both directions for all datasets, but we do not observe a significant difference in performance between the two directions (Supplementary Materials Table 3).

### SCOT’s alternative unsupervised hyperparameter tuning procedure achieves good alignments

We compare the alignment performances in Table 2 when given by SCOT’s alternative tuning procedure guided by the Gromov-Wasserstein distance and MMA-MA’s and UnionCom’s default parameters. SCOT returns nearly the same alignments for simulated data and only marginally worse alignments for real data. In contrast, MMD-MA and UnionCom fail to align some of the simulated and all real datasets with the default parameter values. Therefore, the proposed procedure could guide a user to an alignment close to the optimal result when no orthogonal information is available.

**Table 1:** Alignment performance by label transfer accuracy (*k* = 5).

|  | Sim. 1 | Sim. 2 | Sim. 3 | Syn. RNA-Seq | scGEM | SNAREseq |
| --- | --- | --- | --- | --- | --- | --- |
| **SCOT** | 0.937 | **0.977** | **0.957** | **0.998** | 0.576 | **0.982** |
| **MMD-MA** | 0.89 | 0.783 | 0.947 | 0.706 | **0.588** | 0.942 |
| **UnionCom** | **0.96** | 0.62 | 0.613 | 0.997 | 0.582 | 0.423 |

**Table 2:** Alignment performance by FOSCTTM scores for SCOT chosen by lowest Gromov-Wasserstein distance, default MMD-MA, and default UnionCom for simulated and real datasets.

|  | Sim. 1 | Sim. 2 | Sim. 3 | Syn. RNA-Seq | scGEM | SNAREseq |
| --- | --- | --- | --- | --- | --- | --- |
| **SCOT (GW)** | **0.088** | 0.025 | **0.009** | **0.001** | **0.209** | **0.218** |
| **MMD-MA** | 0.125 | **0.012** | 0.739 | 0.384 | 0.437 | 0.473 |
| **UnionCom** | 0.091 | 0.028 | 0.684 | 0.028 | 0.691 | 0.510 |

### SCOT’s computation speed scales well with the number of samples

We compare SCOT’s running times with the baseline methods for the best performing hyperparameters on the synthetic RNA-seq dataset by varying the number of cells. We run CPU computations on an Intel Xeon e5-2670 with 16GB memory and GPU computations on a single NVIDIA GTX 1080ti with VRAM of 11GB. SCOT’s running time scales similarly to that of MMD-MA, even though SCOT runs on a CPU and MMD-MA runs on a GPU (Figure 4). Both methods scale better than the GPU-based UnionCom implementation.

**Figure 4:**
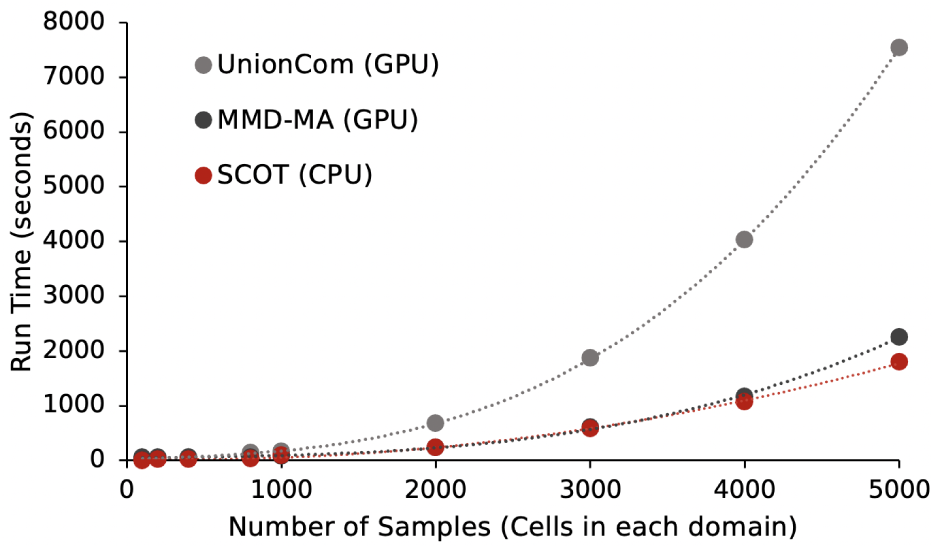
Runtime comparisons with growing sample size. Dotted lines are polynomial trend lines.

## 5 Discussion

We have demonstrated that SCOT, which uses Gromov Wasserstein optimal transport for unsupervised single-cell multi-omics data integration, performs on par with UnionCom and MMD-MA. Our formulation of a coupling matrix based on matching graph distances is somewhat similar to UnionCom’s initial step; however, UnionCom only matches sample-to-sample distances, while Gromov-Wasserstein distance considers the cost of moving pairs of points, enabling our method to better preserve local geometry. Additionally, SCOT performs global alignment of the marginal distributions, which is similar to how MMD-MA uses the MMD term to ensure that the two distributions agree globally in the latent space. We hypothesize that these properties result in SCOT’s state-of-the-art performance. Furthermore, SCOT’s optimization runs in less time and with fewer hyperparameters, and the Gromov-Wasserstein distance can guide the user to choose an alignment when no validation information exists. Therefore, unlike other methods, SCOT easily yields high quality alignments in the fully unsupervised setting.

To visualize and measure alignment, we project data into the same space through barycentric projection, but there are other ways to use the coupling matrix to infer alignment. For example, the coupling matrix could also be used with other dimension reduction methods such as t-SNE (as in UnionCom) to align the manifolds while embedding them both into a new space. Alternatively, depending on the application, a projection may not be required; it may be sufficient to have probabilities relating the samples to one another. Future work will develop effective ways to utilize the coupling matrix and extend our framework to handle more than two alignments at a time.

## Acknowledgments

We are grateful to Yang Lu, Jean-Philippe Vert, and Marco Cuturi for helpful discussion of Gromov-Wasserstein optimal transport.

## Supplementary Materials

### 1 SCOT algorithm

As described in Section 2, SCOT takes in two datasets *X* and *Y* and constructs *k−*NN graphs on each dataset to create the distance matrices *D*_*x*_ and *D*_*y*_. Then, it finds the coupling Γ that minimizes the Gromov-Wasserstein distance. Finally, the coupling matrix is used to project one domain onto the other. In Algorithm 1, we present the full SCOT algorithm, including the Gromov-Wasserstein calculation, which uses the projections proposed in [18].

#### Algorithm 1 Gromov-Wasserstein Alignment

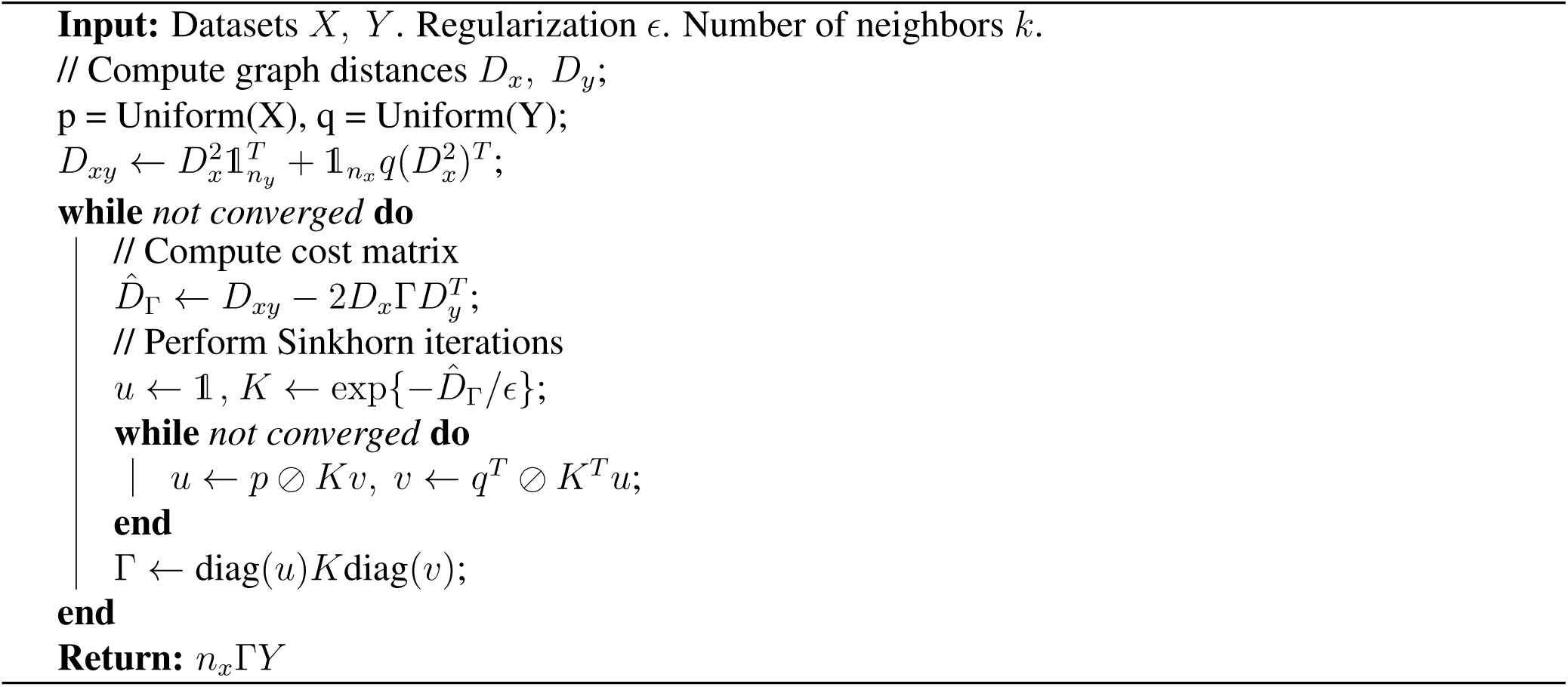

#### 1.1 Unsupervised Hyperparameter Selection Procedure for SCOT

As detailed in Section 2, one way to select SCOT hyperparameters in the absence of correspondence information or validation dataset, is to use the Gromov-Wasserstein distance as a proxy for alignment quality. Here, we present the procedure for carrying this out, where we alternate between the hyperparameters *k* and *E*, and fix one to tune the other:

##### Algorithm 2 Unsupervised hyperparameter search algorithm for SCOT.

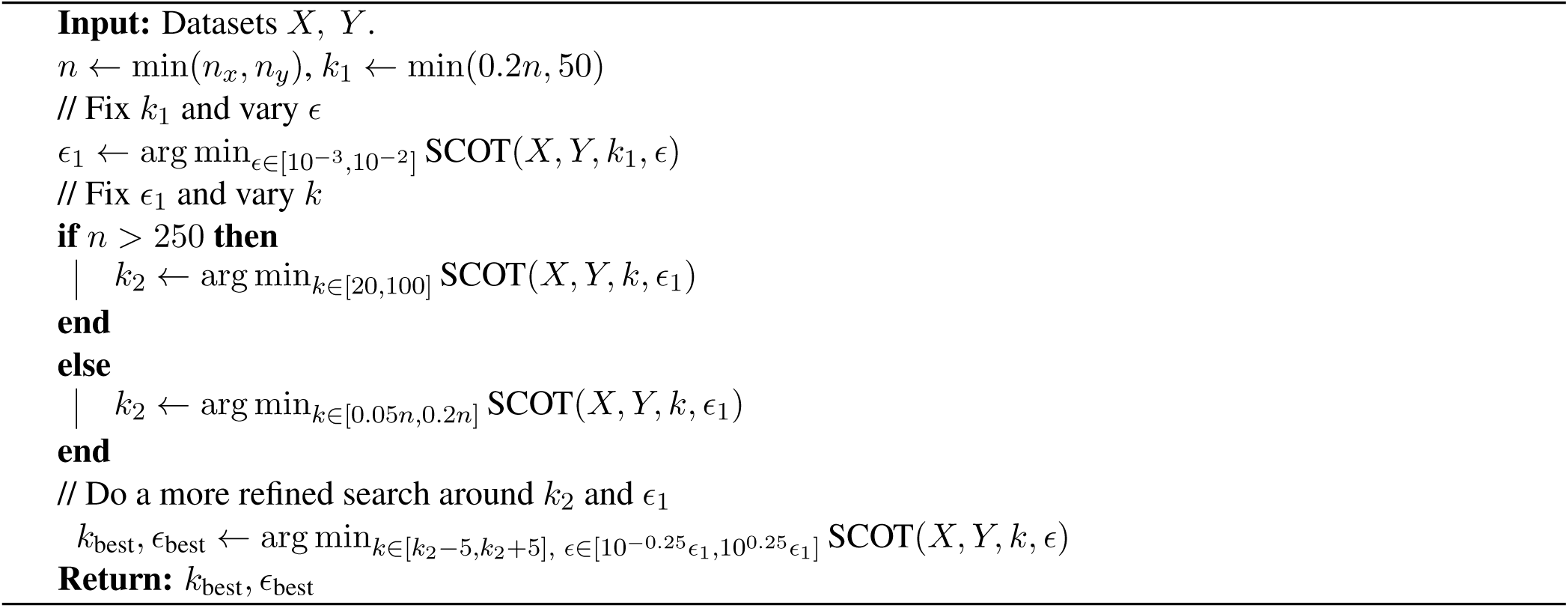

#### 1.2 Visualization of Original Data Sets

In the main text, we display the alignment results performed by SCOT. Here, we visualize the original datasets:

**Figure S1:**
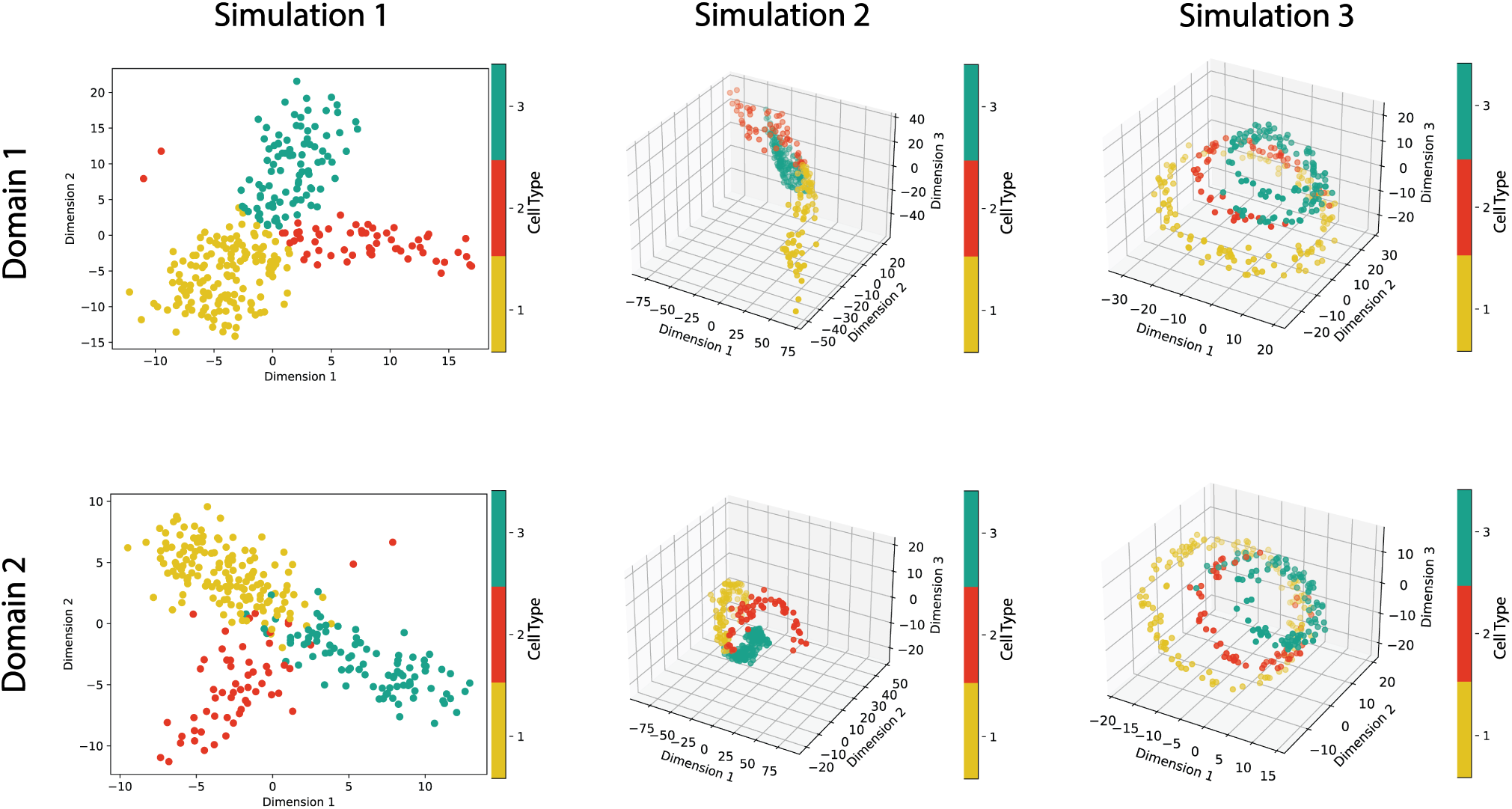
Original simulation data visualized before alignment. Data was generated by Liu et al [8] and retrieved from https://noble.gs.washington.edu/proj/mmd-ma/. Each simulation set has two domains. Their MDS projections in two dimensional and three dimensional space are visualized here. The first set of simulations form a branched tree in two dimensional space (first column); the second set of simulations form Swiss roll in three dimensional space (second column); and lastly, the third set of simulations form a circular frustum.

**Figure S2:**
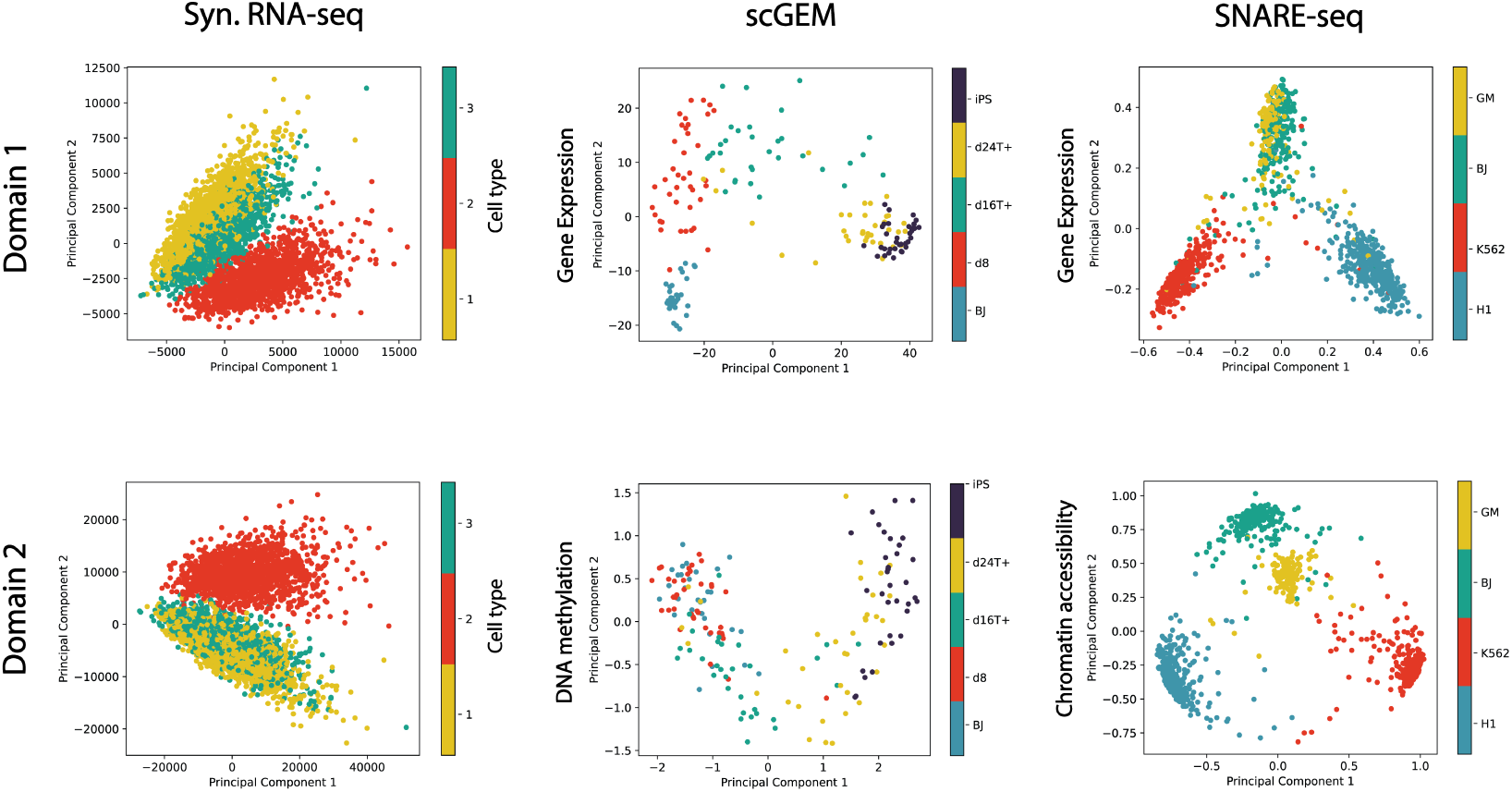
Original synthetic RNA-seq and real world single-cell data visualized before alignment. We use Splatter [21] to generate a count matrix with 5000 cells and 1000 genes from three cell groups. We reduce the dataset to the 5 genes with the highest variances, and then use random matrices to project the data to new dimensions *p*_1_ = 50 and *p*_2_ = 500. Here we visualize the two domains with PCA projections for this dataset as well as the real world single-cell sequnecing datasets

#### 1.3 Barycentric Projections in Both Directions

While MMD-MA and UnionCom project both datasets to a shared low-dimensional space, SCOT projects one dataset onto the other. We project SCOT in both directions for all datasets, but we do not observe a significant difference in performance between the two directions. In Table 3, we present the averaged FOSCTTM values for barycentric projection in both directions (domain 1 projected onto domain 2, as well as domain 2 projected onto domain 1).

**Table 3:**
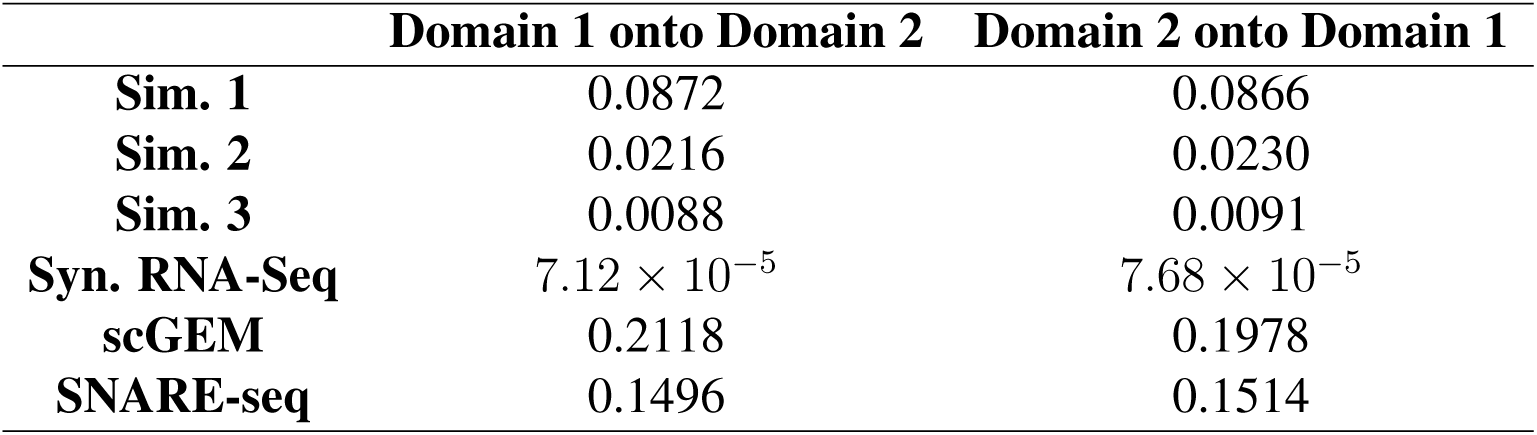
Best mean FOSCTTM for each direction of the barycentric projection for all datasets. The method is robust to the direction of the projection.

**Table 4:**
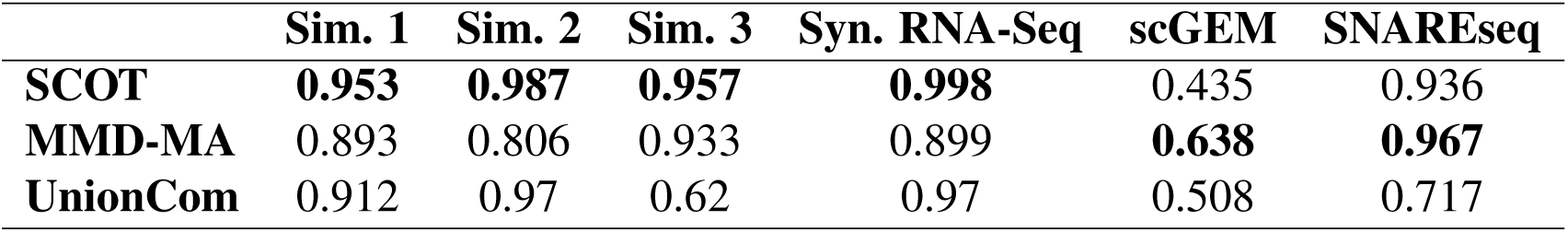
Alignment performance by label transfer accuracy (*k* = 5) for SCOT, MMD-MA, and Union-Com for simulated and real datasets when the second domain is used for training.

**Table 5:** Alignment performance by label transfer accuracy (*k* = 5) when the first domain is used for training for SCOT, MMD-MA, and UnionCom for simulated and real datasets.

|  | **Sim. 1** | **Sim. 2** | **Sim. 3** | **Syn. RNA-Seq** | **scGEM** | **SNAREseq** |
| --- | --- | --- | --- | --- | --- | --- |
| **SCOT** | **0.977** | **0.977** | **0.95** | **0.996** | **0.582** | **0.701** |
| **MMD-MA** | 0.897 | 0.957 | 0.7 | 0.506 | 0.237 | 0.412 |
| **UnionCom** | 0.947 | 0.947 | 0.133 | 0.948 | 0.107 | 0.288 |

**Table 6:**
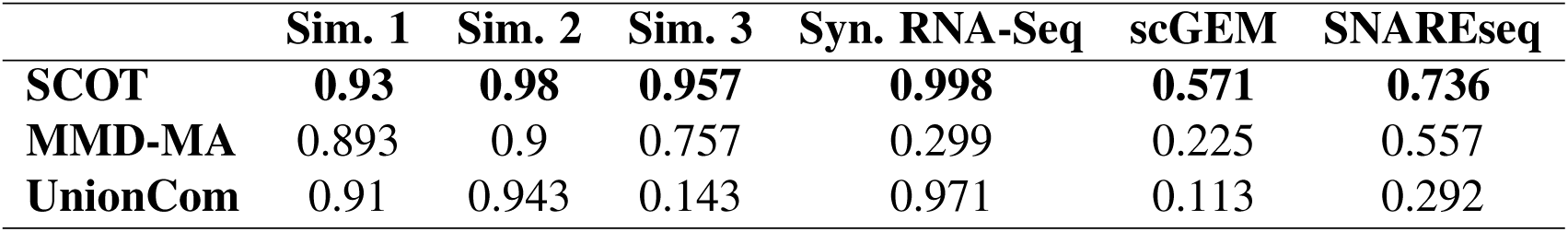
Alignment performance by label transfer accuracy (*k* = 5) when the second domain is used for training for SCOT, MMD-MA, and UnionCom for simulated and real datasets.

#### 1.4 Label Transfer Accuracy with the Second Domain used in Training

In Table 4, we present the label transfer accuracies when the first domain is used as the training set. Here we report the opposite direction.

#### 1.5 Hyperparameter Tuning for SCOT

**Figure S3:**
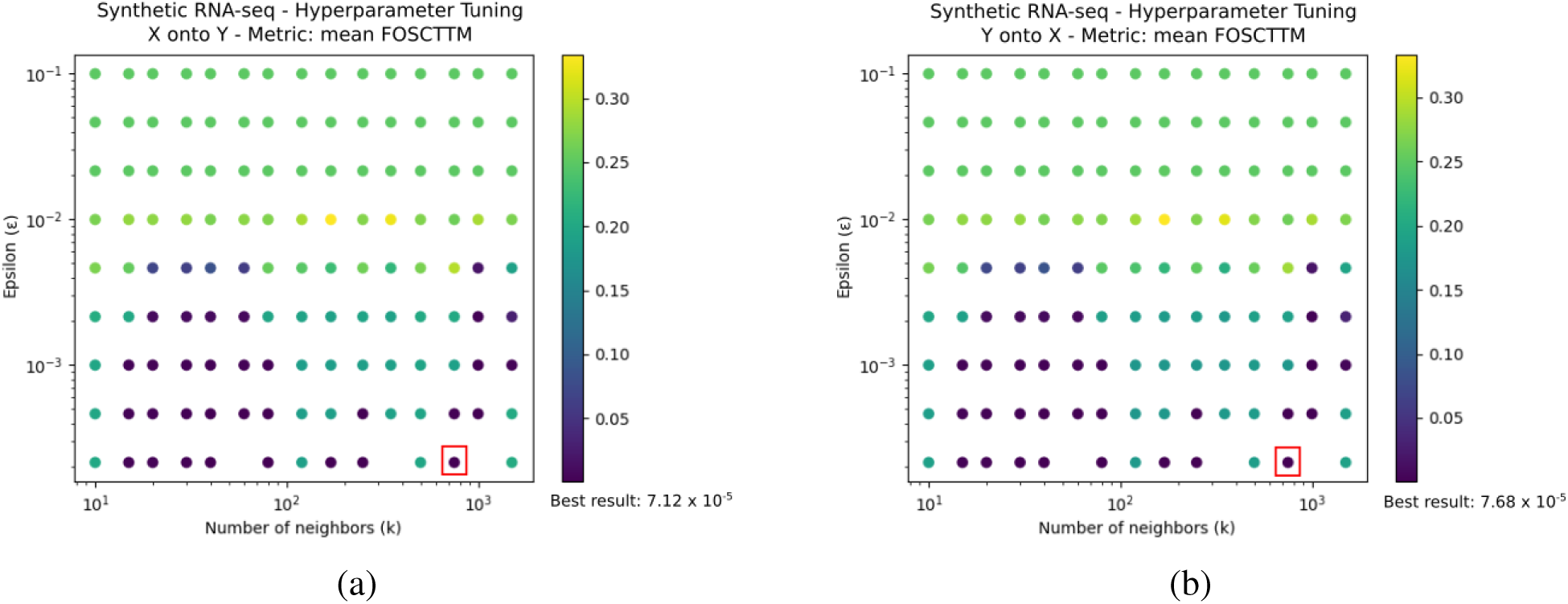
Hyperparameter optimization results for synthetic RNA-seq dataset. Mean FOSCTTM metric was used to assess performance (indicated by color). **(a)** Results when first domain (X) is projected onto second domain (y). **(b)** Results when second domain (y) is projected onto first domain (X). The algorithm is largely robust to the choice of k. For both projections, the best performing hyperparameter setting was *ϵ* = 0.000215, *k* = 750. The hyperparameter combination that yielded the best performance is highlighted with red square. For ease of visualization, a subset of the values are plotted.

**Figure S4:**
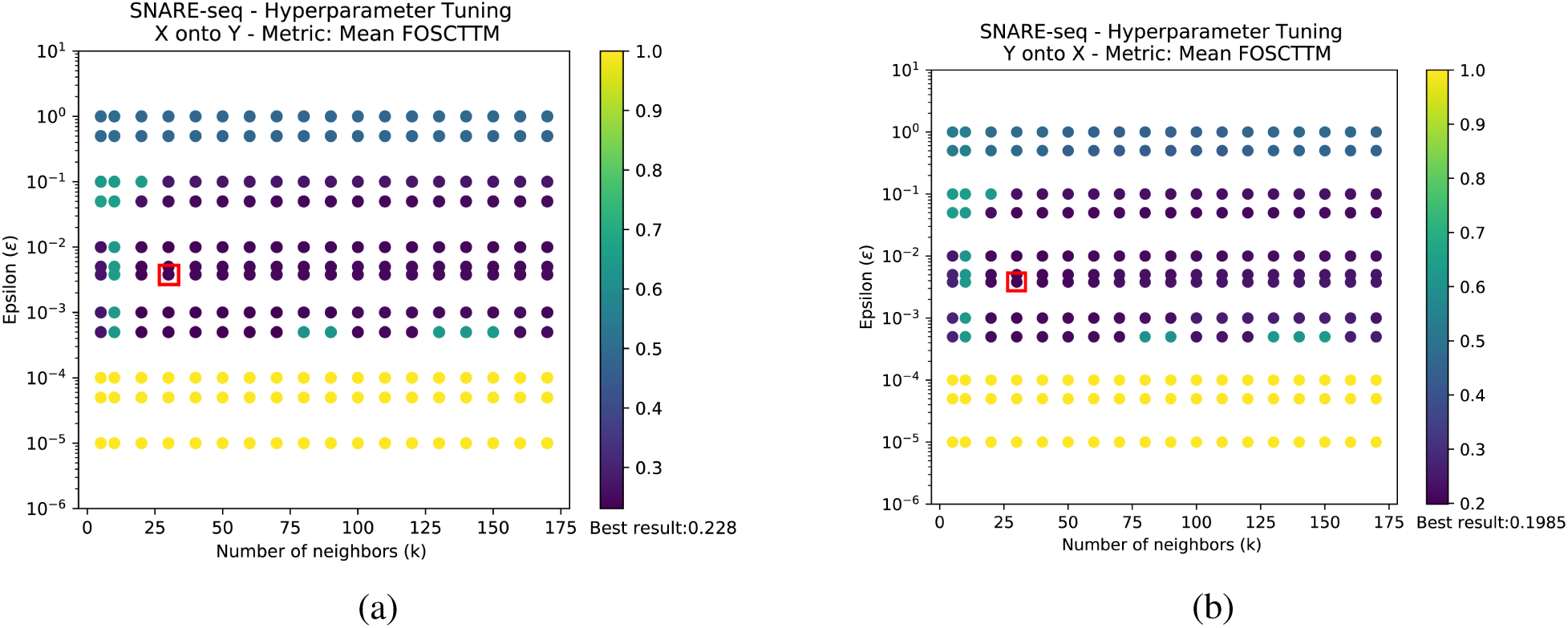
Hyperparameter optimization results for SNARE-seq dataset. Mean FOSCTTM metric was used to assess performance (indicated by color). **(a)** Results when chromatin accessibility domain (X) is projected onto gene expression domain (y). **(b)** Results when expression domain (y) is projected onto chromatin accessibility domain (X). The algorithm is largely robust to the choice of k. For both projections, the best performing hyperparameter setting was *E* = 0.0038, *k* = 30. The hyperparameter combination that yielded the best performance is highlighted with red square. For ease of visualization, a subset of the *ϵ* values are plotted.

#### 1.6 Visualizing the Empirical Relationship between Gromov-Wasserstein Distance and Correspondence in Alignment as Measured by Average FOSCTTM

We observe that lower values of the Gromov-Wasserstein distance tend to correspond to lower average FOSCTTM values. Below, we have plotted the Gromov-Wasserstein values against average FOSCTTM for each dataset over a range of parameter values.

**Figure S5:**
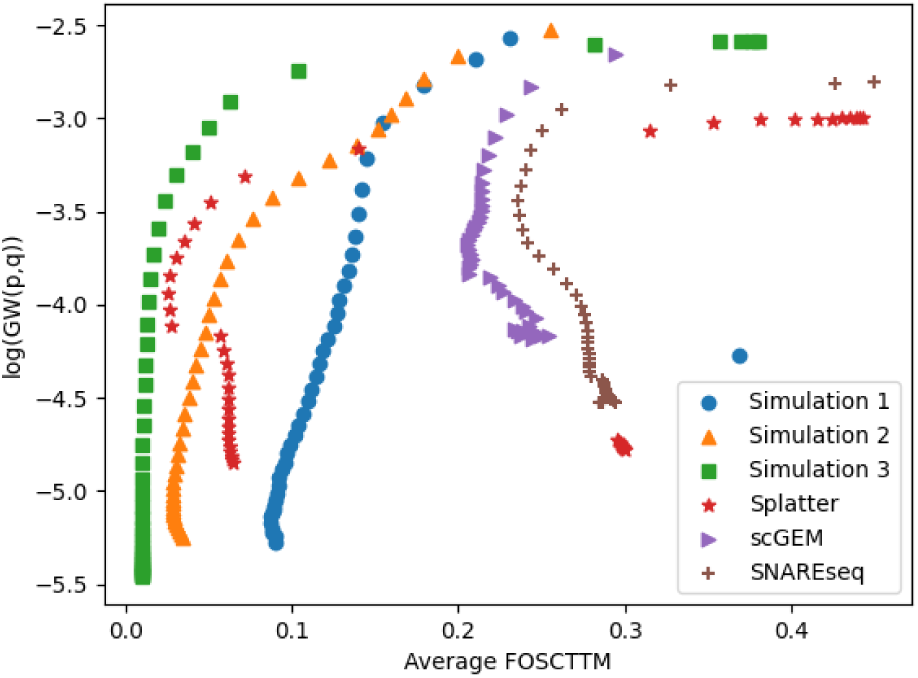
Gromov-Wasserstein distance vs average FOSCTTM values for all datasets with a range of *E* parameter values (*k* fixed at min(50, 0.2*n*_*x*_)).

#### 1.7 Label Transfer Accuracy for Automatic Alignment

In Table 2, we report the average FOSCTTM values for SCOT when chosen by lowest Gromov-Wasserstein distance and default parameters for MMD-MA and UnionCom. In the tables below, we also report the label transfer accuracy scores.

## Footnotes

1 https://noble.gs.washington.edu/proj/mmd-ma/

